## Supplementary material for "Drosophila epidermal cells are intrinsically mechanosensitive and modulate nociceptive behavioral outputs": 14 supplemental figures, 2 supplemental tables, details of statistical analysis, and key resources table

### Figure 1 - Figure supplement 1

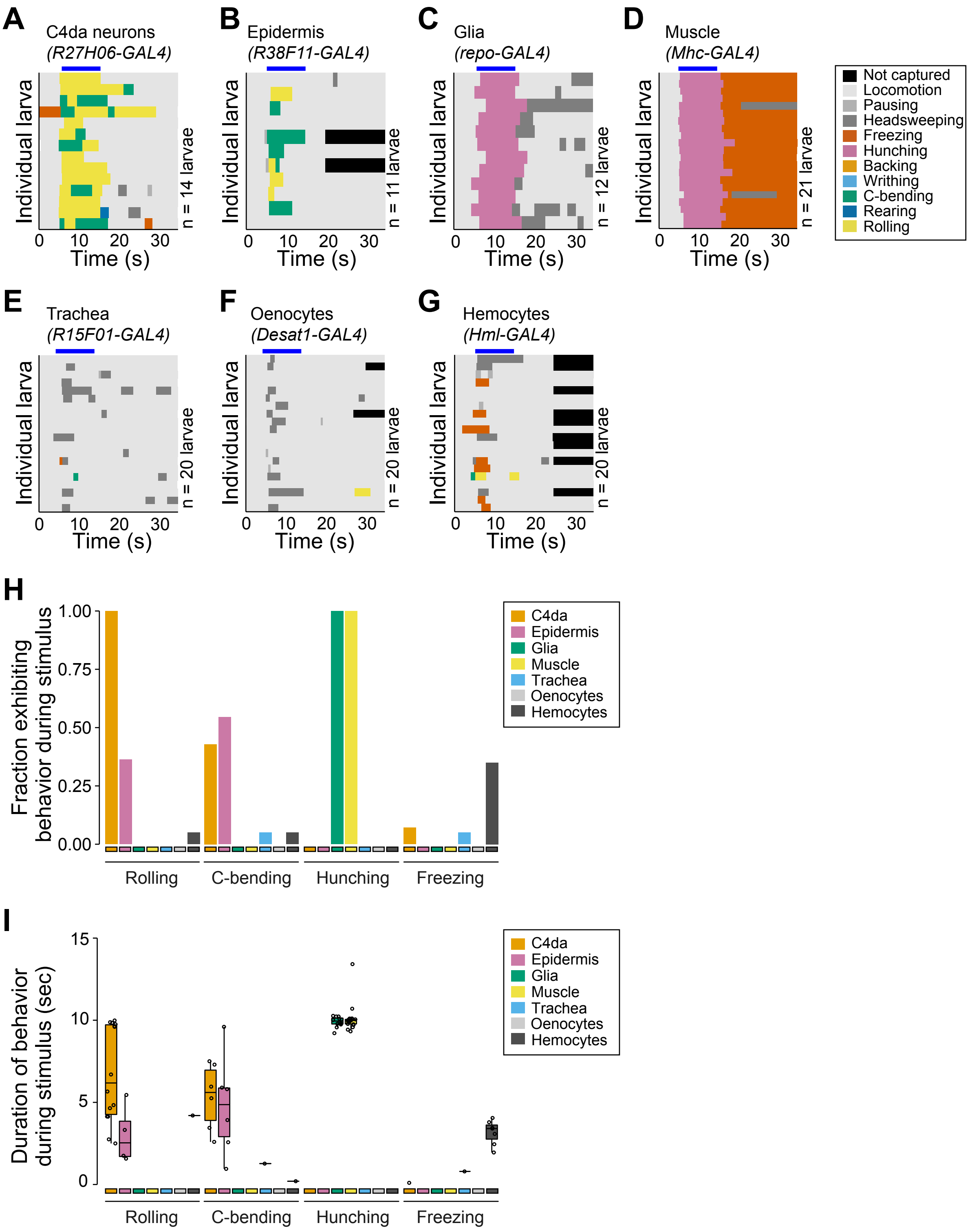

### Figure 1 - Figure supplement 2

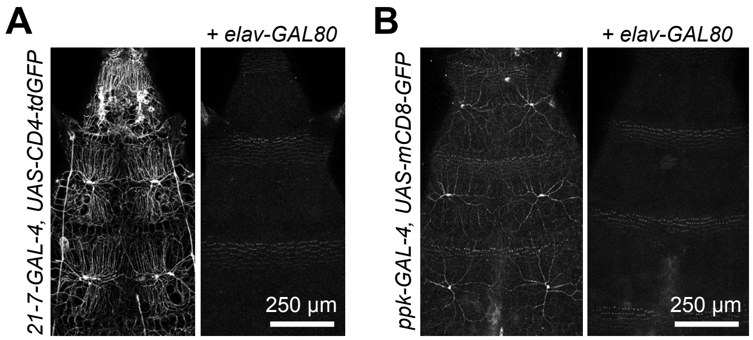

#### Figure 1 - Figure supplement 3

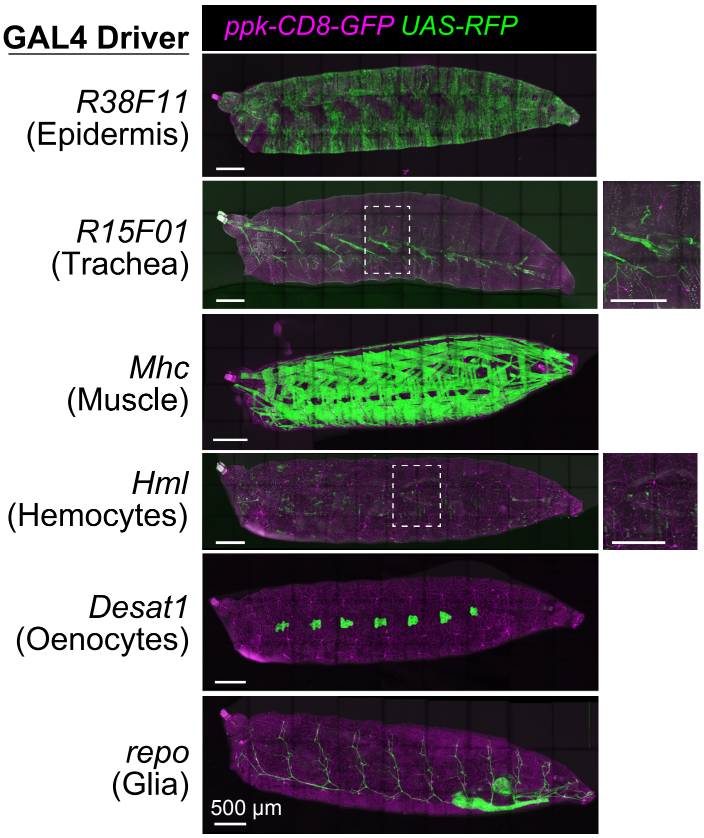

### Figure 1 - Figure supplement 4

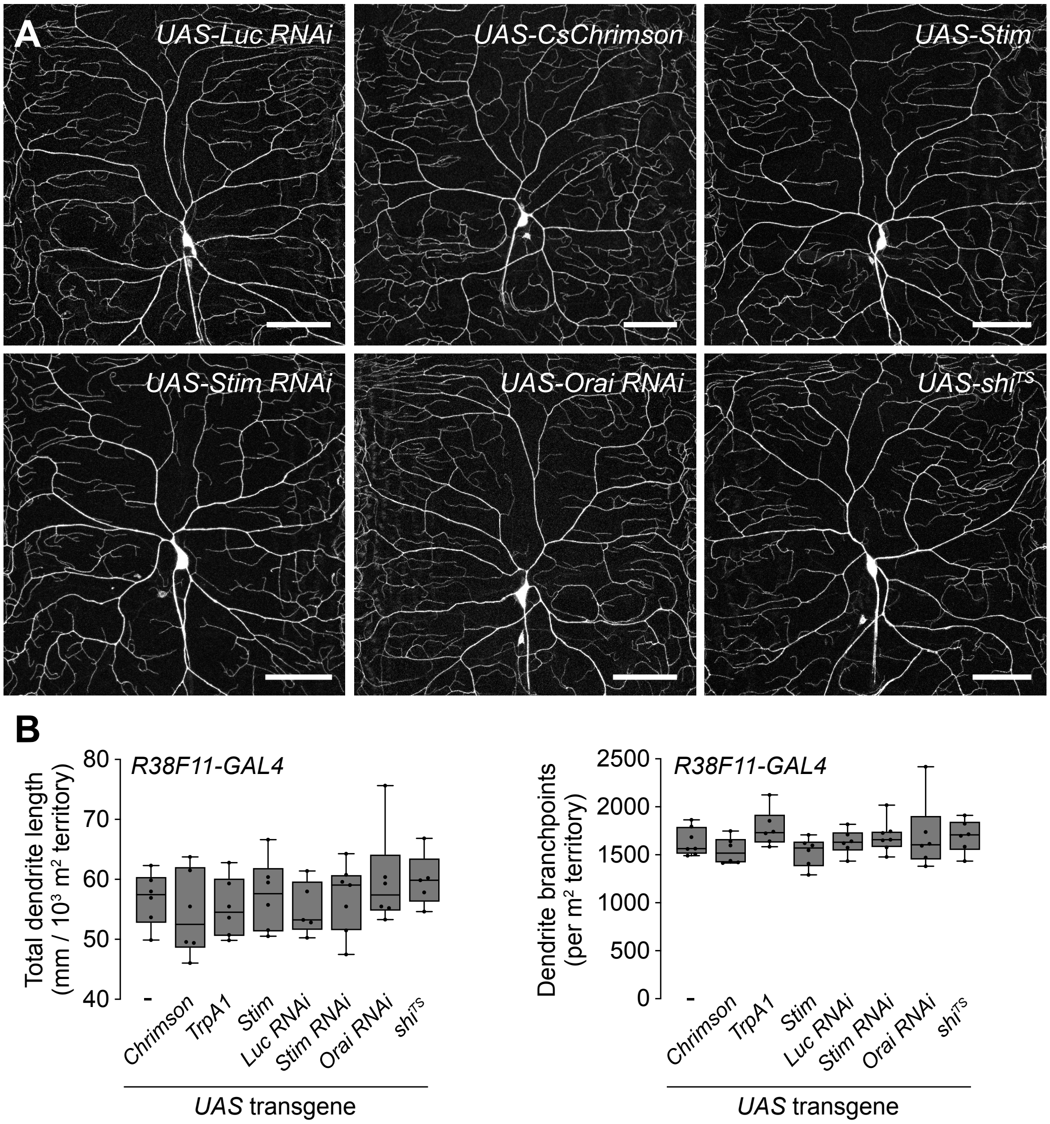

### Figure 1 - Figure supplement 5

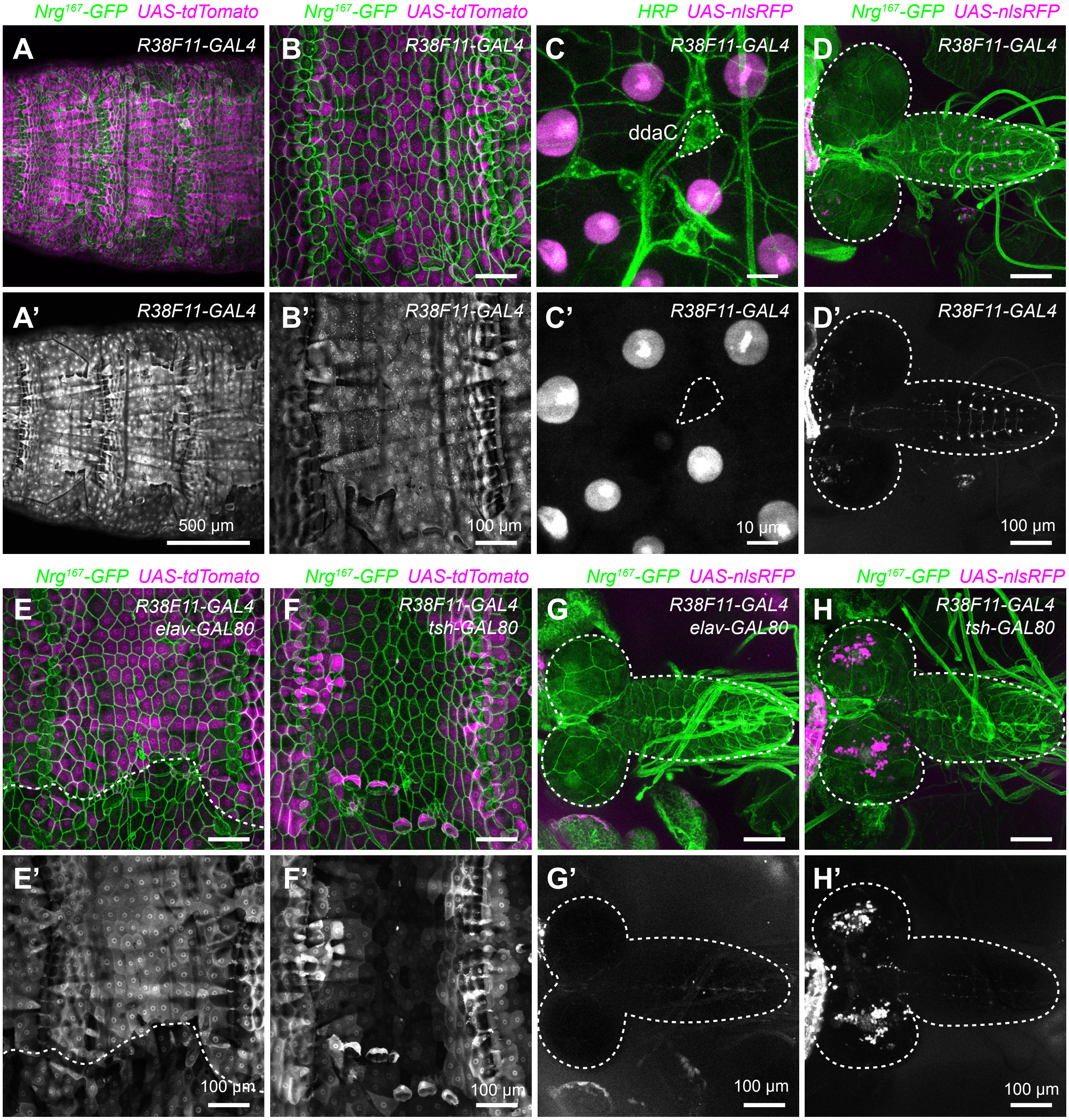

### Figure 1 - Figure supplement 7

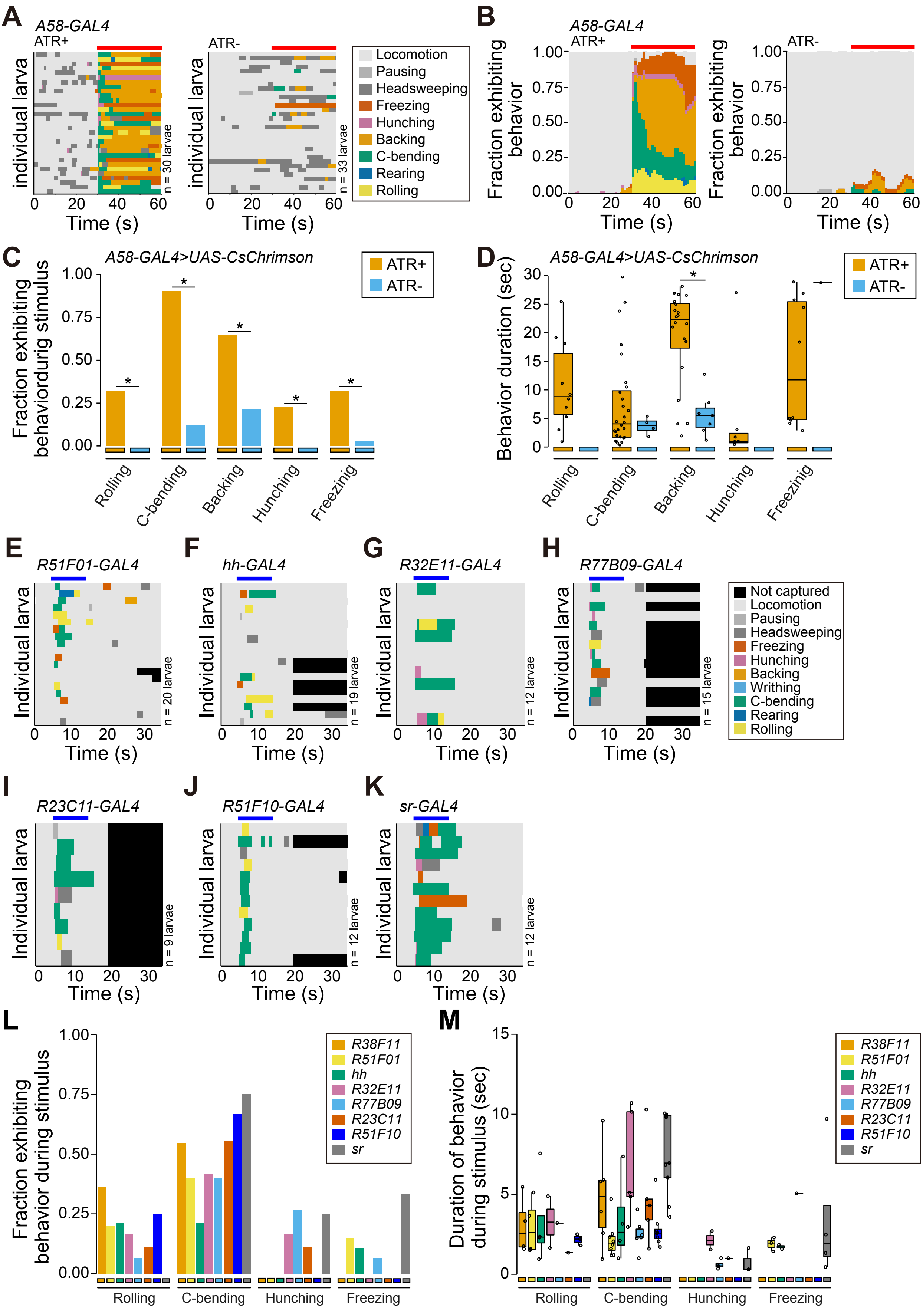

### Figure 1 - Figure supplement 7

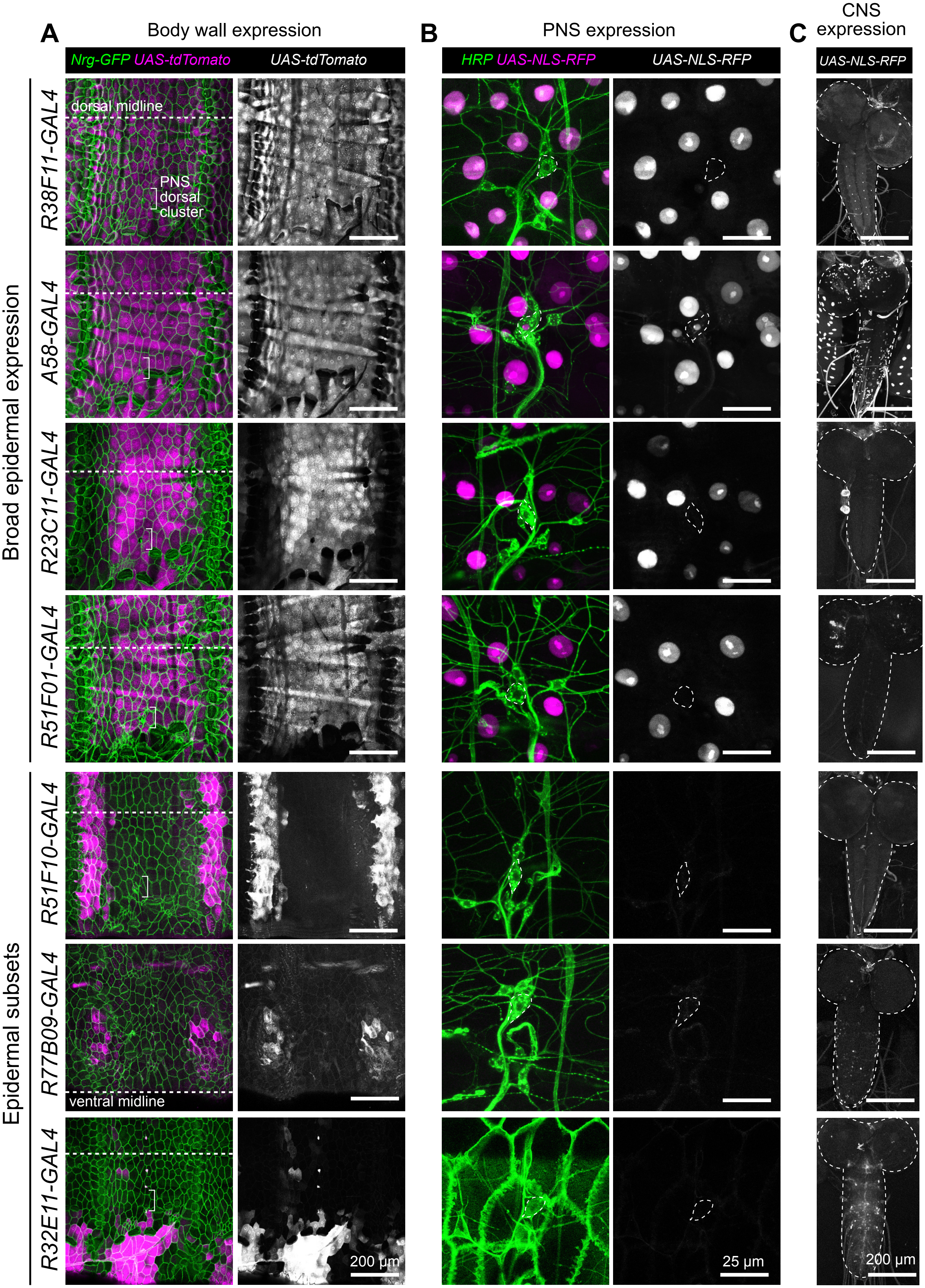

### Figure 2 - Figure supplement 1

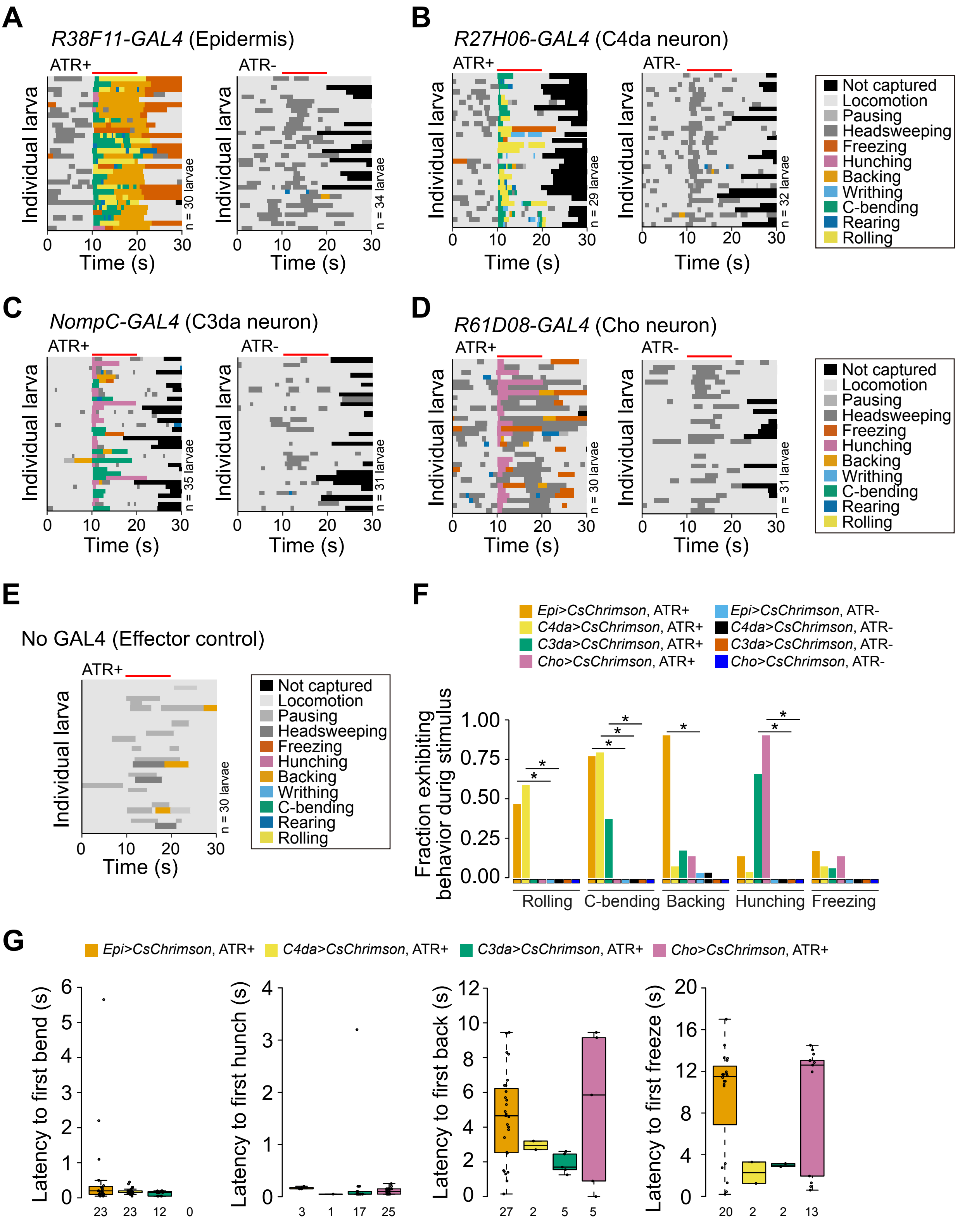

### Figure 3 - Figure supplement 1

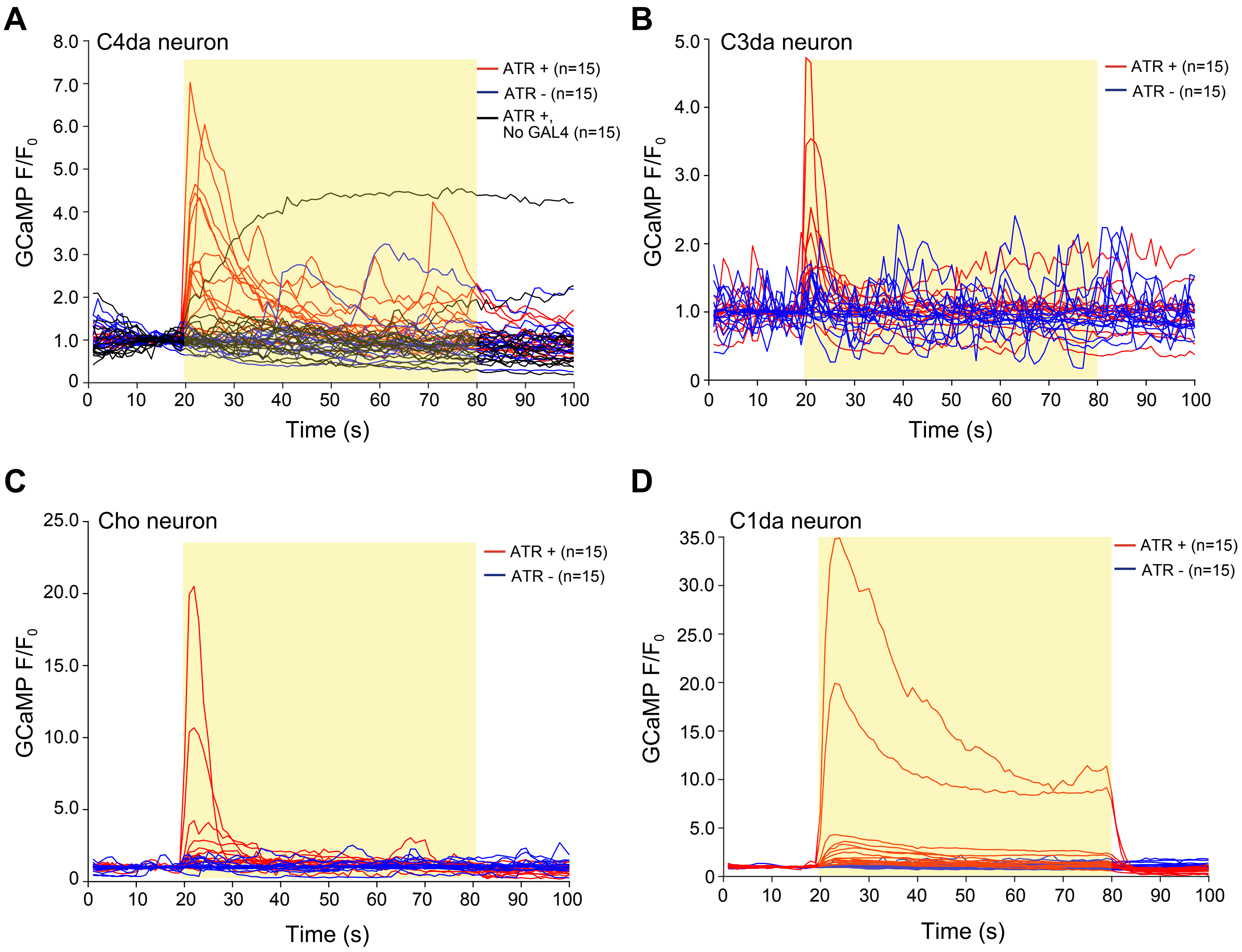

### Figure 3 - Figure supplement 2

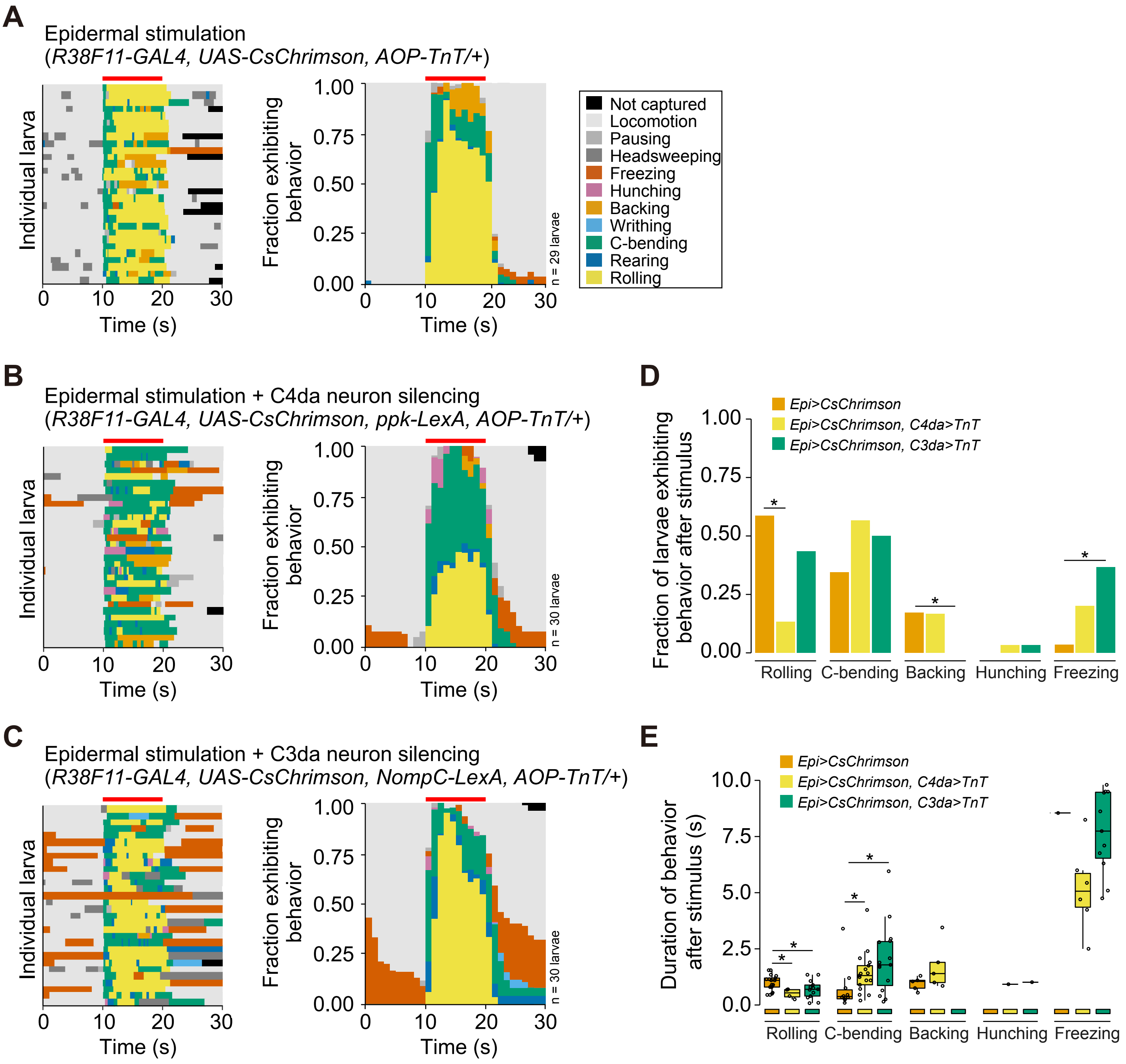

### Figure 4 - Figure supplement 1

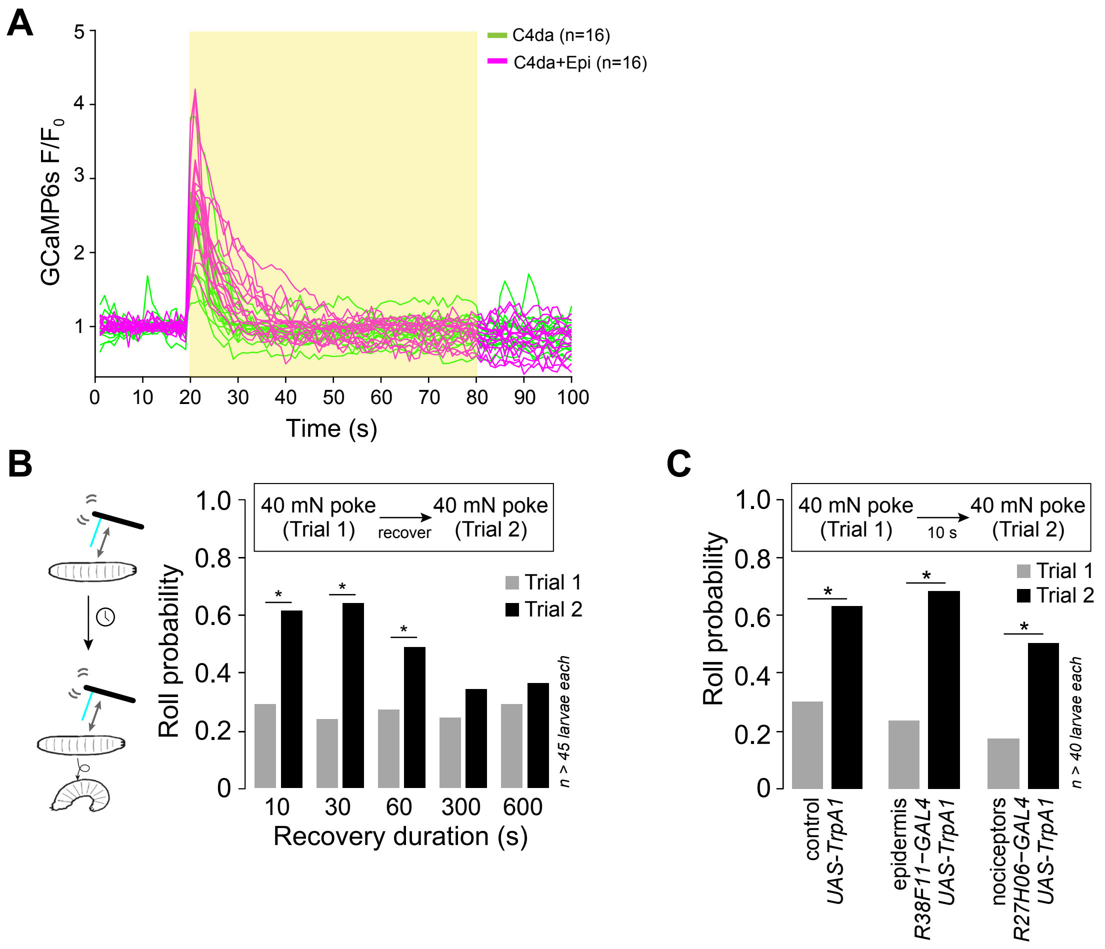

### Figure 5 - Figure supplement 1

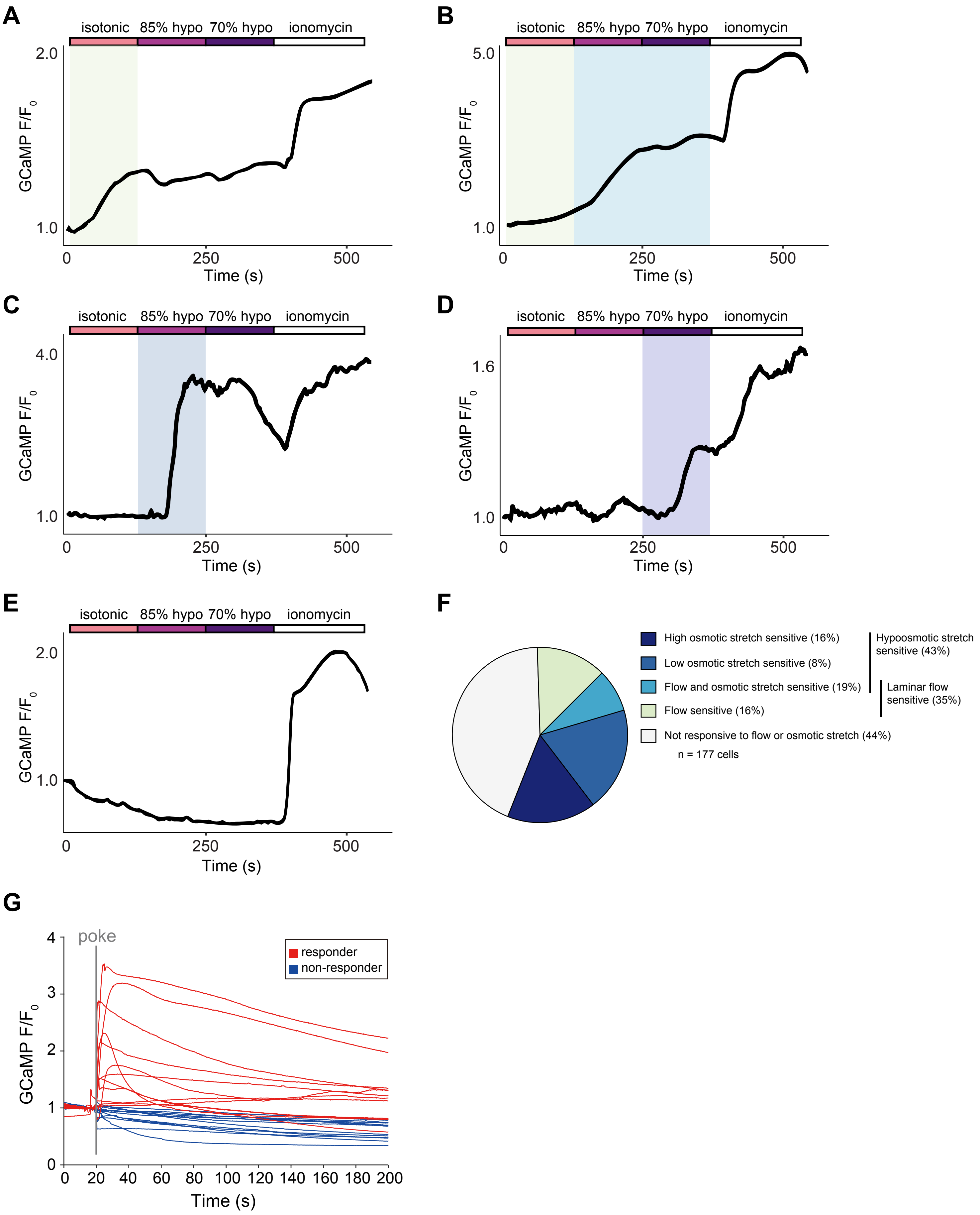

### Figure 6 - Figure supplement 1

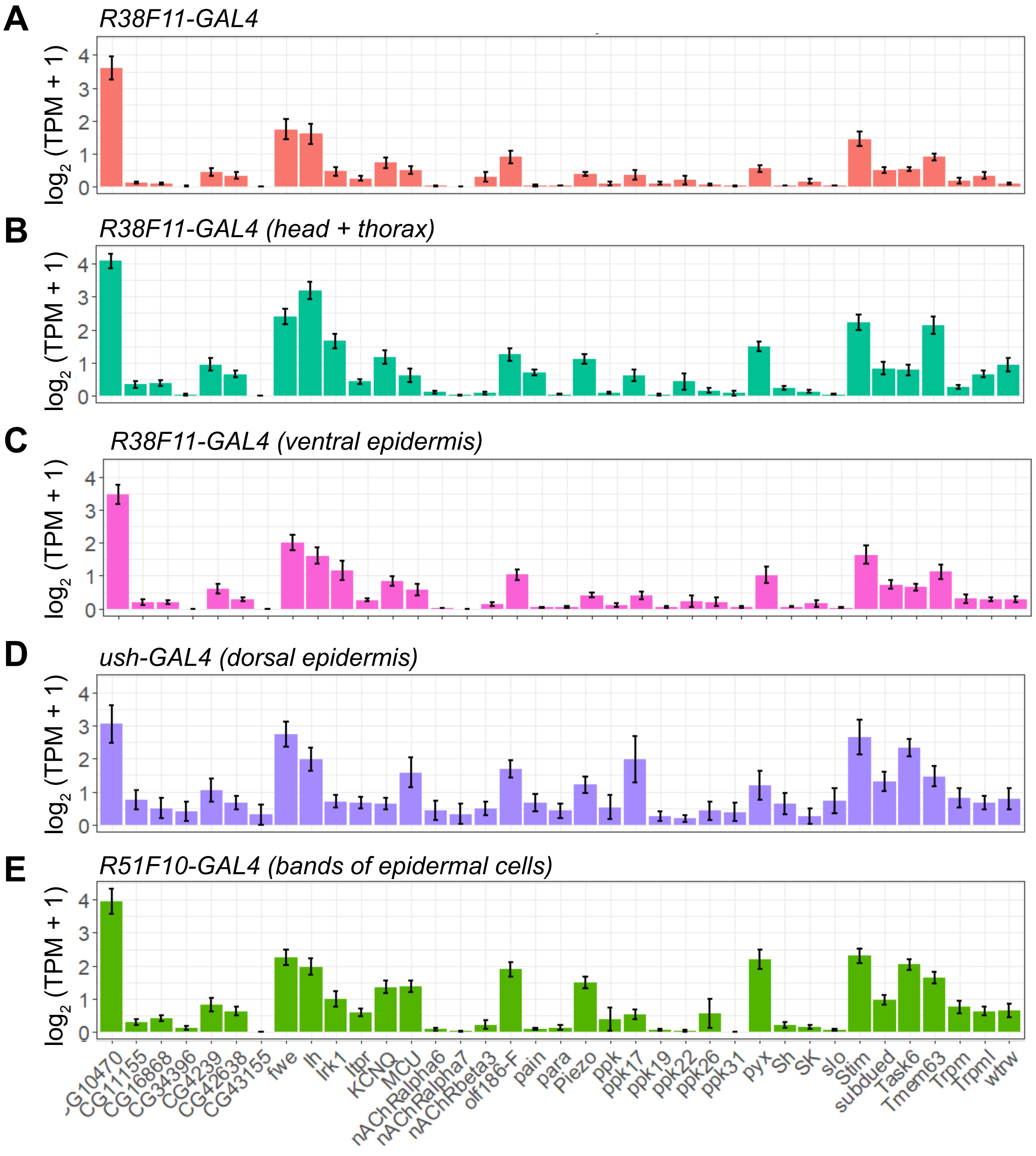

### Figure 6 - Figure supplement 2

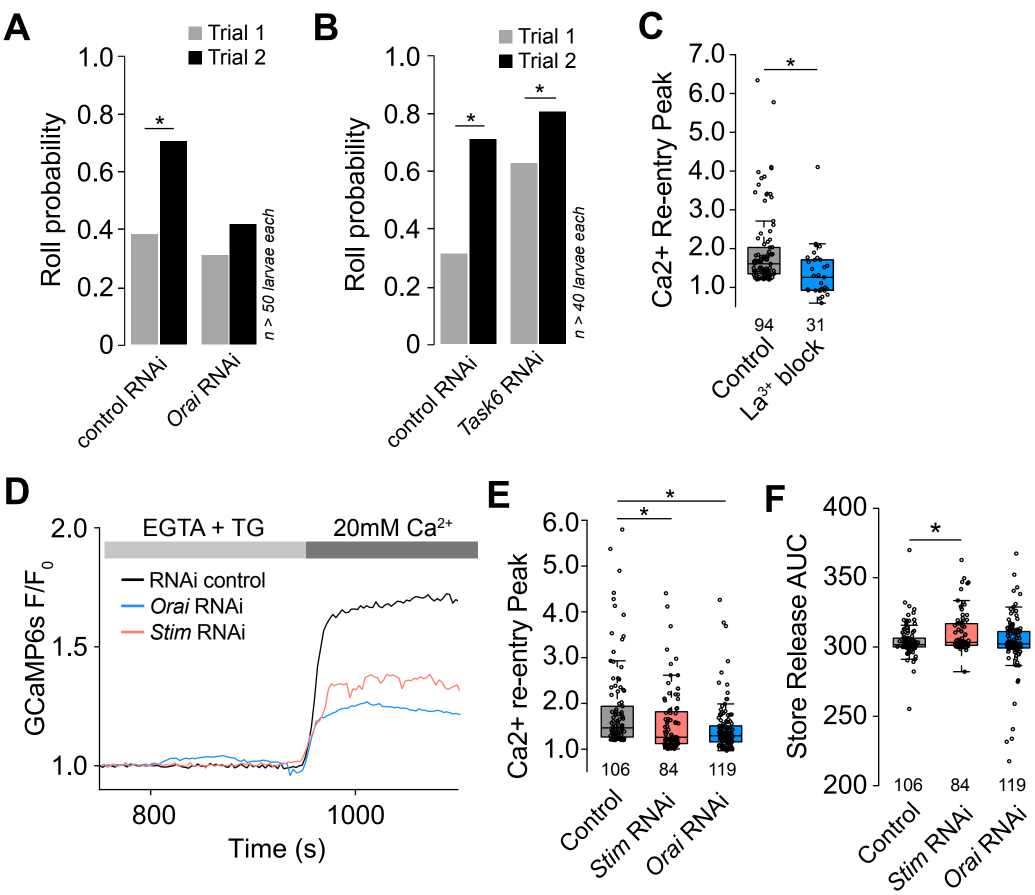

| Key Resources Table | | | | |
|---|---|---|---|---|
| **Reagent type (species) or resource** | **Designation** | **Source or reference** | **Identifiers** | **Additional information** |
| gene (*Drosophila melanogaster*) | Stim | GenBank | FLYB:FBgn 0045073 | |
| gene (*Drosophila melanogaster*) | Orai | GenBank | FLYB:FBgn 0041585 | |
| genetic reagent (*D. melanogaster*) | *A58-GAL4* | Maintained in the Parrish Lab | Flybase_FBal 0181674 | GAL4 driver (epidermis) |
| genetic reagent (*D. melanogaster*) | *Desat1-GAL4* | Bloomington Drosophila Stock Center | BDSC_65405 | GAL4 driver (oenocytes) |
| genetic reagent (*D. melanogaster*) | *GMR15F01-GAL4* | Bloomington Drosophila Stock Center | BDSC_45071 | GAL4 driver (trachea) |
| genetic reagent (*D. melanogaster*) | *GMR23C11-GAL4* | Bloomington Drosophila Stock Center | BDSC_45123 | GAL4 driver (epidermis) |
| genetic reagent (*D. melanogaster*) | *GMR27H06-GAL4* | Bloomington Drosophila Stock Center | BDSC_49440 | GAL4 driver (C4da neurons) |
| genetic reagent (*D. melanogaster*) | *GMR32E11-GAL4* | Bloomington Drosophila Stock Center | BDSC_48109 | GAL4 driver (epidermis) |
| genetic reagent (*D. melanogaster*) | *GMR38F11-GAL4* | Bloomington Drosophila Stock Center | BDSC_50014 | GAL4 driver (epidermis) |

| | | | | |
|---|---|---|---|---|
| genetic reagent (*D. melanogaster*) | *GMR51F01-GAL4* | Bloomington Drosophila Stock Center | BDSC_38787 | GAL4 driver (epidermis) |
| genetic reagent (*D. melanogaster*) | *GMR51F10-GAL4* | Bloomington Drosophila Stock Center | BDSC_38793 | GAL4 driver (epidermis) |
| genetic reagent (*D. melanogaster*) | *GMR61D08-GAL4* | Bloomington Drosophila Stock Center | BDSC_39272 | GAL4 driver (Cho neurons) |
| genetic reagent (*D. melanogaster*) | *GMR77B09-GAL4* | Bloomington Drosophila Stock Center | Flybase_FBti 0138381 | GAL4 driver (epidermis) |
| genetic reagent (*D. melanogaster*) | *hh-GAL4* | Bloomington Drosophila Stock Center | DGGR_11811 7 | GAL4 driver (epidermis) |
| genetic reagent (*D. melanogaster*) | *Hml-GAL4* | Bloomington Drosophila Stock Center | BDSC_6395 | GAL4 driver (hemocytes) |
| genetic reagent (*D. melanogaster*) | *MHC-GAL4* | Bloomington Drosophila Stock Center | BDSC_55132 | GAL4 driver (muscle) |
| genetic reagent (*D. melanogaster*) | *NompC-GAL4* | Bloomington Drosophila Stock Center | BDSC_36361 | GAL4 driver (C3da neurons) |
| genetic reagent (*D. melanogaster*) | *Piezo-GAL4* | Bloomington Drosophila Stock Center | BDSC_78335 | GAL4 driver (Piezo-expressing cells) |
| genetic reagent (*D. melanogaster*) | *ppk-GAL4* | Bloomington Drosophila Stock Center | BDSC_32079 | GAL4 driver (C4da neurons) |

| | | | | |
|---|---|---|---|---|
| genetic reagent (*D. melanogaster*) | *21-7-GAL4* | Maintained in the Parrish Lab | Flybase_FBti 0131369 | GAL4 driver (MD neurons) |
| genetic reagent (*D. melanogaster*) | *repo-GAL4* | Bloomington Drosophila Stock Center | BDSC_7415 | GAL4 driver (glia) |
| genetic reagent (*D. melanogaster*) | *sr-GAL4* | Bloomington Drosophila Stock Center | BDSC_26663 | GAL4 driver (apodemes) |
| genetic reagent (*D. melanogaster*) | *ppk-LexA* | Maintained in the Parrish Lab | Flybase_FBtp 0125814 | LEXA driver (C4da neurons) |
| genetic reagent (*D. melanogaster*) | *NompC-LexA* | Bloomington Drosophila Stock Center | BDSC_52241 | LEXA driver (C3da neurons) |
| genetic reagent (*D. melanogaster*) | *elav-GAL80* | Maintained in the Parrish Lab | Flybase_FBtp 0079702 | GAL80 (pan-neuronal) |
| genetic reagent (*D. melanogaster*) | *tsh-GAL80* | Bloomington Drosophila Stock Center | Flybase_FBti 0114123 | GAL80 (VNC) |
| genetic reagent (*D. melanogaster*) | *ush-GAL4* | Bloomington Drosophila Stock Center | BDSC_36524 | GAL4 driver (epidermis) |
| genetic reagent (*D. melanogaster*) | *Nrg167GFP* | Bloomington Drosophila Stock Center | BDSC_6844 | Reporter (epidermal cell junctions) |
| genetic reagent (*D. melanogaster*) | *UAS-tdTomato* | Bloomington Drosophila Stock Center | BDSC_36328 | Reporter (RFP) |

| | | | | |
|---|---|---|---|---|
| genetic reagent (*D. melanogaster*) | *UAS-RedStinger* | Bloomington Drosophila Stock Center | BDSC_8546 | Reporter (NLS-RFP) |
| genetic reagent (*D. melanogaster*) | *UAS0GCaMP6s* | Bloomington Drosophila Stock Center | BDSC_42749 | Reporter (Calcium indicator) |
| genetic reagent (*D. melanogaster*) | *UAS-GCaMP6s* | Bloomington Drosophila Stock Center | BDSC_42746 | Reporter (Calcium indicator) |
| genetic reagent (*D. melanogaster*) | *AOP-GCaMP6s* | Bloomington Drosophila Stock Center | BDSC_44273 | Reporter (Calcium indicator) |
| genetic reagent (*D. melanogaster*) | *AOP-TNT* | Maintained in the Parrish Lab | Flybase_FBtp 0144631 | Neuronal silencing (Tetanus Toxin) |
| genetic reagent (*D. melanogaster*) | *UAS-shi-ts* | Bloomington Drosophila Stock Center | BDSC_44222 | Inducible inhibition of endocytosis |
| genetic reagent (*D. melanogaster*) | *UAS-CsChrimson* | Bloomington Drosophila Stock Center | BDSC_55135 | Inducible Cation Channel |
| genetic reagent (*D. melanogaster*) | *UAS-CsChrimson* | Bloomington Drosophila Stock Center | BDSC_55136 | Inducible Cation Channel |
| genetic reagent (*D. melanogaster*) | *UAS-TRPA1* | Bloomington Drosophila Stock Center | BDSC_26263 | Inducible Cation Channel |
| genetic reagent (*D. melanogaster*) | *UAS-GtACR* | Bloomington Drosophila Stock Center | BDSC_92983 | Inducible Anion Channel |

| | | | | |
|---|---|---|---|---|
| genetic reagent (*D. melanogaster*) | *UAS-luciferase RNAi* | Bloomington Drosophila Stock Center | BDSC_31603 | RNAi transgene |
| genetic reagent (*D. melanogaster*) | *UAS-RFP-RNAi* | Bloomington Drosophila Stock Center | BDSC_67852 | RNAi transgene |
| genetic reagent (*D. melanogaster*) | *UAS-fwe-RNAi* | Bloomington Drosophila Stock Center | BDSC_27323 | RNAi transgene |
| genetic reagent (*D. melanogaster*) | *UAS-Ih-RNAi* | Bloomington Drosophila Stock Center | BDSC_29574 | RNAi transgene |
| genetic reagent (*D. melanogaster*) | *UAS-Ih-RNAi* | Bloomington Drosophila Stock Center | BDSC_58089 | RNAi transgene |
| genetic reagent (*D. melanogaster*) | *UAS-Irk1-RNAi* | Bloomington Drosophila Stock Center | BDSC_42644 | RNAi transgene |
| genetic reagent (*D. melanogaster*) | *UAS-KCNQ-RNAi* | Bloomington Drosophila Stock Center | BDSC_80446 | RNAi transgene |
| genetic reagent (*D. melanogaster*) | *UAS-MCU-RNAi* | Bloomington Drosophila Stock Center | BDSC_67897 | RNAi transgene |
| genetic reagent (*D. melanogaster*) | *UAS-orai-RNAi* | Bloomington Drosophila Stock Center | BDSC_53333 | RNAi transgene |
| genetic reagent (*D. melanogaster*) | *UAS-orai-RNAi* | Vienna Drosophila Resource Center | VDRC_12221 | RNAi transgene |

| | | | | |
|---|---|---|---|---|
| genetic reagent (*D. melanogaster*) | *UAS-pain-RNAi* | Bloomington Drosophila Stock Center | BDSC_51835 | RNAi transgene |
| genetic reagent (*D. melanogaster*) | *UAS-piezo-RNAi* | Vienna Drosophila Resource Center | FlyBase_FBst 0457216 | RNAi transgene |
| genetic reagent (*D. melanogaster*) | *UAS-piezo-RNAi* | Kyoto Drosophila Stock Center | Flybase_FBtp 0071516 | RNAi transgene |
| genetic reagent (*D. melanogaster*) | *UAS-pyx-RNAi* | Bloomington Drosophila Stock Center | BDSC_31297 | RNAi transgene |
| genetic reagent (*D. melanogaster*) | *UAS-stim-RNAi* | Bloomington Drosophila Stock Center | BDSC_41759 | RNAi transgene |
| genetic reagent (*D. melanogaster*) | *UAS-stim-RNAi* | Vienna Drosophila Resource Center | FlyBase_FBst 0478081 | RNAi transgene |
| genetic reagent (*D. melanogaster*) | *UAS-subdued-RNAi* | Vienna Drosophila Resource Center | FlyBase_FBst 0480747 | RNAi transgene |
| genetic reagent (*D. melanogaster*) | *UAS-Task6-RNAi* | Bloomington Drosophila Stock Center | BDSC_28016 | RNAi transgene |
| genetic reagent (*D. melanogaster*) | *UAS-Task6-RNAi* | Vienna Drosophila Resource Center | FlyBase_FBst 0471360 | RNAi transgene |
| genetic reagent (*D. melanogaster*) | *UAS-Tmem63-RNAi* | Vienna Drosophila Resource Center | FlyBase_FBst 0470667 | RNAi transgene |

| | | | | |
|---|---|---|---|---|
| genetic reagent (*D. melanogaster*) | *UAS-TMCO1-RNAi* | Bloomington Drosophila Stock Center | BDSC_42896 | RNAi transgene |
| genetic reagent (*D. melanogaster*) | *UAS-TMCO1-RNAi* | Bloomington Drosophila Stock Center | BDSC_55909 | RNAi transgene |
| genetic reagent (*D. melanogaster*) | *UAS-TrpA1-RNAi* | Bloomington Drosophila Stock Center | BDSC_66905 | RNAi transgene |
| genetic reagent (*D. melanogaster*) | *UAS-Trpm-RNAi* | Bloomington Drosophila Stock Center | BDSC_31291 | RNAi transgene |
| genetic reagent (*D. melanogaster*) | *UAS-Trpml-RNAi* | Bloomington Drosophila Stock Center | BDSC_31294 | RNAi transgene |
| genetic reagent (*D. melanogaster*) | *UAS-wtrw-RNAi* | Bloomington Drosophila Stock Center | BDSC_51563 | RNAi transgene |
| cell line (*Homo-sapiens*) | Immortalized Human Keratinocytes (HaCaT cells) | Cytion | Cat. #: 300493 | |
| peptide, recombinant protein | Collagenase type I | Thermo Fisher | Cat. #: 17-100-017 | |
| chemical compound, drug | Fura-2AM | Thermo Fisher | Cat. #: F1221 | |
| chemical compound, drug | Pluronic F-127 | Thermo Fisher | Cat. #: P3000MP | |

| | | | | |
|---|---|---|---|---|
| chemical compound, drug | All-trans retinal | Millipore Sigma | Cat. #: R2500 | |
| chemical compound, drug | ionomycin | Thermo Fisher | Cat. #: I24222 | |
| chemical compound, drug | Lanthanum chloride | Millipore Sigma | Cat. #: 211605 | 500 nM |
| software, algorithm | SPSS | SPSS | RRID:SCR_002865 | |
| software, algorithm | Matlab | Mathworks | RRID:SCR_001622 | |
| software, algorithm | ImageJ | ImageJ | RRID:SCR_003070 | |
| software, algorithm | CASAVA | Illumina, http://www.illumina.com/software/genome_analyzer_software.ilmn | | |
| software, algorithm | FasQC | Babraham bioinformatics, https://www.bioinformatics.babraham.ac.uk/ | | |
| software, algorithm | Cutadapt v2.4 | (Martin, 2011) | | |
| software, algorithm | Kallisto version 0.46.0 | (Bray et al, 2016) | | |

#### Supplementary File 1: Details of Statistical Analysis

**Figure 1C**: Fisher's exact test with BH correction

| *UAS-TRPA1* groups (25 C vs 32 C) | p-value | q-value | Significance (q<0.05) |
|---|---|---|---|
| Control (no *GAL4* driver) | 0.2424 | 2.424000e-01 | F |
| *R27H06-GAL4* | 2.2e-16 | 3.666667e-16 | T |
| *R38F11-GAL4* | 2.2e-16 | 3.666667e-16 | T |
| *R38F11-GAL4 + elav-GAL80* | 2.2e-16 | 3.666667e-16 | T |
| *R38F11-GAL4 + tshGAL80* | 4.111e-15 | 5.138750e-15 | T |

**Figure 1S4**: ANOVA with post-hoc Tukey's test

| Dendrite length | p-value | Significance |
|---|---|---|
| *R38 vs. UAS-luc RNAi* | >0.9999 | F |
| *R38 vs. UAS-CsChrimson* | 0.9659 | F |
| *R38 vs. UAS-TRPA1* | 0.9982 | F |
| *R38 vs. UAS-orai RNAi* | 0.9009 | F |
| *R38 vs. UAS-Stim RNAi* | 0.9219 | F |
| *R38 vs. UAS-Stim* | >0.9999 | F |
| *R38 vs. UAS-shi^TS* | 0.9974 | F |

| Dendrite branchpoints | p-value | Significance |
|---|---|---|
| *R38 vs. UAS-luc RNAi* | 0.9945 | F |
| *R38 vs. UAS-CsChrimson* | 0.9269 | F |
| *R38 vs. UAS-TRPA1* | 0.6859 | F |
| *R38 vs. UAS-orai RNAi* | 0.9896 | F |
| *R38 vs. UAS-Stim RNAi* | 0.9949 | F |
| *R38 vs. UAS-Stim* | 0.8654 | F |
| *R38 vs. UAS-shi^TS* | >0.9999 | F |

**Figure 1S6C**: Fisher's exact test

| | p-value | Significance |
|---|---|---|
| rolling | 0.0002928 | T |
| c-bend | 2.035e-10 | T |
| backing | 0.0008424 | T |
| hunch | 0.004233 | T |
| freeze | 0.002342 | T |

**Figure 1S6D**: Wilcoxon rank sum test

| | p-value | Significance |
|---|---|---|
| rolling | Cannot compare | |
| c-bend | 0.8907 | F |
| backing | 0.00127 | T |
| hunch | Cannot compare | |
| freeze | Cannot compare | |

**Figure 2F:** Wilcoxon Rank-Sum test

| | p-value | Significance |
|---|---|---|
| C4da vs Epi | 0.0067 | T |

**Figure 2G:** Kruskal-Wallis test followed by Wilcoxon rank sum test with BH correction

| | p-value | q-value | Significance (q<0.05) |
|---|---|---|---|

| | | | |
|---|---|---|---|
| Kruskal-Wallis test, rolling | N/A (only epi + C4da have values) | | |
| rolling C4da vs Epi | 0.03124 | | T |
| Kruskal-Wallis test, c-bend | 0.0001652 | | T |
| c-bend C4da vs Epi | 0.001468 | 0.002936 | T |
| c-bend C3da vs Epi | 0.3237 | 0.323700 | F |

**Figure 2H** Fisher's exact test with a BH correction

| | p-value | q-value | Significance (q<0.05) |
|---|---|---|---|
| rolling Epi ATR+ vs ATR- | 0.04313 | 0.09102 | F |
| rolling C4da ATR+ vs ATR- | 0.04551 | 0.09102 | F |
| rolling C3da ATR+ vs ATR- | 1 | 1 | F |
| rolling Cho ATR+ vs ATR | 1 | 1 | F |
| c-bend Epi ATR+ vs ATR- | 0.4687 | 0.9508 | F |
| c-bend C4da ATR+ vs ATR- | 0.4754 | 0.9508 | F |
| c-bend C3da ATR+ vs ATR | 1 | 1 | F |
| c-bend Cho ATR+ vs ATR | 1 | 1 | F |
| back Epi ATR+ vs ATR- | 0.00000000000004556 | 0.00000000000018224 | T |
| back C4da ATR+ vs ATR | 1 | 1 | F |
| back C3da ATR+ vs ATR | 1 | 1 | F |
| back Cho ATR+ vs ATR- | 0.1128 | 0.2256 | F |
| hunch Epi ATR+ vs ATR | 1 | 1 | F |
| hunch C4da ATR+ vs ATR | 1 | 1 | F |
| hunch C3da ATR+ vs ATR- | 1 | 1 | F |
| hunch Cho ATR+ vs ATR- | 0.4918 | 1 | F |
| freeze Epi ATR+ vs ATR- | 0.00000008682 | 0.00000034728 | T |
| freeze C4da ATR+ vs ATR- | 0.4754 | 0.6338667 | F |
| freeze C3da ATR+ vs ATR- | 1 | 1 | F |
| freeze Cho ATR+ vs ATR- | 0.000825 | 0.00165 | T |

**Figure 2S1E:** Fisher's exact test with BH correction

| | p-value | q-value | Significance (q<0.05) |
|---|---|---|---|
| rolling Epi ATR+ vs ATR- | 3.039e-06 | 6.0780e-06 | T |
| rolling C4da ATR+ vs ATR- | 9.667e-08 | 3.8668e-07 | T |
| rolling C3da ATR+ vs ATR- | 1 | 1.0000e+00 | F |
| rolling Cho ATR+ vs ATR- | 1 | 1.0000e+00 | F |
| c-bend Epi ATR+ vs ATR- | 1.388e-11 | 2.776e-11 | T |
| c-bend C4da ATR+ vs ATR- | 1.265e-11 | 2.776e-11 | T |
| c-bend C3da ATR+ vs ATR- | 0.00192 | 2.560e-03 | T |
| c-bend Cho ATR+ vs ATR- | 1 | 1.000e+00 | F |
| back Epi ATR+ vs ATR- | 1.238e-13 | 4.952000e-13 | T |
| back C4da ATR+ vs ATR- | 0.6003 | 6.003000e-01 | F |
| back C3da ATR+ vs ATR- | 0.02597 | 5.194000e-02 | F |
| back Cho ATR+ vs ATR- | 0.05251 | 7.001333e-02 | F |
| hunch Epi ATR+ vs ATR- | 0.04313 | 5.750667e-02 | F |
| hunch C4da ATR+ vs ATR- | 0.4754 | 4.754000e-01 | F |
| hunch C3da ATR+ vs ATR- | 1.03e-06 | 2.060000e-06 | T |
| hunch Cho ATR+ vs ATR- | 2.571e-14 | 1.028400e-13 | T |
| freeze Epi ATR+ vs ATR- | 0.01869 | 0.0747600 | F |
| freeze C4da ATR+ vs ATR- | 0.2219 | 0.2958667 | F |
| freeze C3da ATR+ vs ATR- | 0.5353 | 0.5353000 | F |
| freeze Cho ATR+ vs ATR- | 0.05251 | 0.1050200 | F |

**Figure 3E** Fisher's exact test with BH correction

| | p-value | q-value | Significance (q<0.05) |
|---|---|---|---|
| rolling control vs *C4da>TNT* | 0.001051 | 0.002102 | T |
| rolling control vs *C3da>TNT* | 0.1945 | 0.194500 | F |
| c-bend control vs *C4da>TNT* | 1 | 1.00000 | F |
| c-bend control vs *C3da>TNT* | 0.05219 | 0.10438 | F |
| back control vs *C4da>TNT* | 0.7787 | 0.778700 | F |
| back control vs *C3da>TNT* | 0.005579 | 0.011158 | T |
| hunch control vs *C4da>TNT* | 0.02372 | 0.04744 | T |
| hunch control vs *C3da>TNT* | 0.1124 | 0.11240 | F |
| freeze control vs *C4da>TNT* | 0.3533 | 0.3533 | F |
| freeze control vs *C3da>TNT* | 0.1945 | 0.3533 | F |

**Figure 3F** Kruskal-Wallis test followed by Wilcoxon rank sum test with BH correction

| | p-value | q-value | Significance (q<0.05) |
|---|---|---|---|
| Kruskal-Wallis test, rolling | 0.0004424 | | |
| rolling control vs *C4da>TNT* | 0.001122 | 0.002244 | T |
| rolling control vs *C3da>TNT* | 0.7007 | 0.700700 | F |
| Kruskal-Wallis test, c-bend | 0.001948 | | |
| c-bend control vs *C4da>TNT* | 0.0004278 | 0.0008556 | T |
| c-bend control vs *C3da>TNT* | 0.433 | 0.4330000 | F |
| Kruskal-Wallis test, back | 0.4203 | | |
| back control vs *C4da>TNT* | N/A | | |
| back control vs *C3da>TNT* | N/A | | |
| hunch control vs *C4da>TNT* | No Hunch in control | | |
| hunch control vs C3da TNT | No Hunch in control<br>(p=0.4127 C4vsC3) | | |
| Kruskal-Wallis test, freeze | 0.3808 | | |
| freeze control vs *C4da>TNT* | N/A | | |
| freeze control vs *C3da>TNT* | N/A | | |

**Figure 3S2 D**: Fisher's exact test with BH correction

| | p-value | q-value | Significance (q<0.05) |
|---|---|---|---|
| Control vs C4da silence, roll | 0.0003682 | 0.0007364 | T |
| Control vs C3da silence, roll | 0.3015 | 0.3015000 | F |
| Control vs C4da silence, c-bend | 0.1188 | 0.2376 | F |
| Control vs C3da silence, c-bend | 0.2949 | 0.2949 | F |
| Control vs C4da silence, backward | 1 | 1.00000 | F |
| Control vs C3da silence, backward | 0.02372 | 0.04744 | T |
| Control vs C4da silence, hunch | 1 | 1 | F |
| Control vs C3da silence, hunch | 1 | 1 | F |
| Control vs C4da silence, freeze | 0.1028 | 0.102800 | F |
| Control vs C3da silence, freeze | 0.002466 | 0.004932 | T |

**Figure 3S2 E**: Kruskal-Wallis test followed by Wilcoxon rank sum test

| | p-value | q-value | Significance |
|---|---|---|---|
| Kruskal-Wallis test, Roll | 0.02471 | | T |
| Control vs C4da silence Roll | 0.01604 | 0.03208 | T |
| Control vs C3da silence Roll | 0.04658 | 0.04658 | T |
| Kruskal-Wallis test, c-bend | 0.01971 | | T |
| Control vs C4da silence c-bend | 0.008927 | 0.017854 | T |
| Control vs C3da silence c-bend | 0.02334 | 0.023340 | T |

| | | | |
|---|---|---|---|
| Control vs C4da silence back | 0.2222 | | F |
| Control vs C3da silence back | N/A | | |
| Control vs C3da silence back | N/A | | |
| Control vs C4da silence hunch | N/A | | |
| Control vs C3da silence hunch | N/A | | |
| Control vs C4da silene freeze | N/A | | |
| Control vs C3da silence freeze | N/A | | |

**Figure 4B**: Wilcoxon Rank-Sum test

| | p-value | Significance |
|---|---|---|
| Fmax/F0 C4da vs C4da +Epi | 0.0288 | T |

**Figure 4C**: Wilcoxon Rank Sum test

| | p-value | q-value | Significance (q<0.05) |
|---|---|---|---|
| C4da vs C4da+Epi | 0.03184 | | T |

**Figure 4D**: Wilcoxon Rank-Sum test

| | p-value | Significance |
|---|---|---|
| C4da vs C4da+Epi | 0.03277 | T |

**Figure 4F**: Fisher's exact test with BH correction

| | p-value | q-value | Significance (q<0.05) |
|---|---|---|---|
| C4da vs Epi | 0.0003136 | 3.741e-04 | T |
| C4da vs Epi+C4da | 0.7541 | 3.741e-04 | T |
| Epi vs Epi+C4da | 0.000005162 | 2.997e-11 | T |

**Figure 4G**: Kruskal-Wallis test followed by Wilcoxon rank sum test with BH correction

| | p-value | q-value | Significance (q<0.05) |
|---|---|---|---|
| Kruskal-Wallis test | 0.000000001544 | | |
| C4da vs Epi | 0.7541 | 0.7541 | F |
| C4da vs Epi+C4da | 0.000000000001072 | 0.000000000003216 | T |
| Epi vs Epi+C4da | 0.000005162 | 0.000007743 | T |

**Figure 4H:** Fisher's exact test with BH correction

| | p-value | q-value | Significance (q<0.05) |
|---|---|---|---|
| <5 C4da vs Epi | 0.1894 | 1.894e-01 | F |
| <5 C4da vs C4da+Epi | 3.646e-11 | 5.469e-11 | T |
| <5 Epi vs C4da+Epi | 2.571e-14 | 7.713e-14 | T |
| 5-9 C4da vs Epi | 0.674 | 0.674 | F |
| 5-9 C4da vs C4da+Epi | 0.356 | 0.674 | F |
| 5-9 Epi vs C4da+Epi | 0.6124 | 0.674 | F |
| 10 -14 C4da vs Epi | 0.6144 | 0.9216 | F |
| 10 -14 C4da vs C4da+Epi | 1 | 1.0000 | F |
| 10 -14 Epi vs C4da+Epi | 0.6124 | 0.9216 | F |
| 15 -19 C4da vs Epi | 1 | 1.0000 | F |
| 15-19 C4da vs C4da+Epi | 0.1896 | 0.2844 | F |
| 15 -19 Epi vs C4da+Epi | 0.1128 | 0.2844 | F |
| >20 C4da vs Epi | 1 | 1.0000 | F |
| >20 C4da vs C4da+Epi | 5.377e-11 | 1.6131e-10 | T |
| >20 Epi vs C4da+Epi | 2.644e-10 | 3.9660e-10 | T |

**Figure 4I**: Kruskal-Wallis test followed by Wilcoxon rank sum test with BH correction

| | p-value | q-value | Significance (q<0.05) |
|---|---|---|---|
| Kruskal-Wallis test | 5.993e-15 | | |
| C4da vs Epi | 0.0009667 | 9.6670e-04 | T |
| C4da vs Epi+C4da | 4.97e-15 | 7.4550e-15 | T |
| Epi vs Epi+C4da | 5.028e-16 | 1.5084e-15 | T |

**Figure 4J**: Kruskal-Wallis test followed by Wilcoxon rank sum test with BH correction

| | p-value | q-value | Significance (q<0.05) |
|---|---|---|---|
| Kruskal-Wallis test, rolling | 4.024e-09 | | T |
| roll C4da vs Epi | 0.1937 | 1.9370e-01 | F |
| roll C4da vs C4da+Epi | 9.024e-12 | 2.7072e-11 | T |
| roll Epi vs C4da + Epi | 5.506e-05 | 8.2590e-05 | T |
| Kruskal-Wallis test, c-bend | 0.06175 | | F |
| c-bend C4da vs Epi | N/A | | |
| c-bend C4da vs C4da+Epi | N/A | | |
| c-bend Epi vs C4da +Epi | N/A | | |
| Kruskal-Wallis test, backing | 9.145e-06 | | T |
| back C4da vs Epi | 9.574e-07 | 2.8722e-06 | T |
| back C4da vs C4da+Epi | 0.02662 | 2.6620e-02 | T |
| back Epi vs C4da+Epi | 0.001987 | 2.9805e-03 | T |
| Kruskal-Wallis test, hunching | 0.006437 | | T |
| hunch C4da vs Epi | 0.2706 | 0.270600 | F |
| hunch C4da vs C4da+Epi | 0.001252 | 0.003756 | T |
| hunch Epi vs C4da+Epi | 0.1836 | 0.270600 | F |
| Kruskal-Wallis test, freezing | 0.4857 | | F |
| freeze C4da vs Epi | N/A | | |
| freeze C4da vs C4da+Epi | N/A | | |
| freeze Epi vs C4da+Epi | N/A | | |

**Figure 4K**: Fisher's exact test with a BH correction

| *Epi-GAL4, UAS-CsChrimson* treatment groups | p-value | q-value | Significance |
|---|---|---|---|
| 20 mN ATR- vs ATR+ Epi>CsChrimson | 0.003234 | 0.0097020 | T |
| 50 mN ATR- vs ATR+ Epi>CsChrimson | 0.008797 | 0.0131955 | T |
| 50mN ATR- vs ATR+ CsChrimson | 0.7639 | 0.7639000 | F |

**Figure 4L**: Fisher's exact test with BH correction

| | p-value | q-value | Significance (q<0.05) |
|---|---|---|---|
| No *GAL4* 25 vs 32 | 0.6798 | 0.7806000 | F |
| *Epi-GAL4* 25 vs 32 | 0.0005454 | 0.0021816 | T |
| *27H06-GAL4* 25 vs 32 | 0.7806 | 0.7806000 | F |
| *ppk-GAL4* 25 vs 32 | 0.7735 | 0.7806000 | F |

**Figure 4M**: Fisher's exact test with BH correction

| *UAS-TRPA1* comparisons | p-value | q-value | Significance (q<0.05) |
|---|---|---|---|
| 10sec control vs 10sec *Epi-GAL4* | 0.006108 | 0.01018000 | T |
| 30sec control vs 30sec *Epi-GAL4* | 0.001265 | 0.00316250 | T |
| 60sec control vs 60sec *Epi-GAL4* | 0.00007129 | 0.00035645 | T |
| 300sec control vs 300sec *Epi-GAL4* | 0.07531 | 0.09413750 | F |
| 600sec control vs 600sec *Epi-GAL4* | 0.4351 | 0.43510000 | F |

**Figure 4N**: Curve fitting

| | fit curve | Decay time constant |
|---|---|---|
| Mechano (Fig 4S1B) | f(x) = 0.4139*exp( -0.002994*x) | 334.001336 |
| Epidermal activation (Fig 4L) | f(x) = 0.47*exp(-0.002964*x) | 337.3819163 |

**Figure 4S1B**: Fisher's exact test with BH correction

| | p-value | q-value | Significance (q<0.05) |
|---|---|---|---|
| 10sec $1^{st}$ vs $2^{nd}$ | 0.0007546 | 0.0018865 | T |
| 30sec $1^{st}$ vs $2^{nd}$ | 0.000006865 | 0.000034325 | T |
| 60sec $1^{st}$ vs $2^{nd}$ | 0.008544 | 0.01424 | T |
| 300sec $1^{st}$ vs $2^{nd}$ | 0.131 | 0.16375 | F |
| 600sec $1^{st}$ vs $2^{nd}$ | 0.3217 | 0.3217 | F |

**Figure 4S1C**: Fisher's exact test with BH correction

| | p-value | q-value | Significance (q<0.1) |
|---|---|---|---|
| Control $1^{st}$ vs $2^{nd}$ | 0.006746 | 0.00674600 | T |
| *Epi-GAL4* $1^{st}$ vs $2^{nd}$ | 6.336e-05 | 0.00019008 | T |
| *Nociceptor-GAL4* $1^{st}$ vs $2^{nd}$ | 0.005186 | 0.00674600 | T |

**Figure 6B** Fisher's exact test with BH correction

| | p-value | q-value | Significance (q<0.1) |
|---|---|---|---|
| Control $1^{st}$ vs $2^{nd}$ | 4.94E-05 | 0.0002472 | T |
| *Orai* RNAi $1^{st}$ vs $2^{nd}$ | 0.1329 | 0.3236667 | F |
| *Stim* RNAi $1^{st}$ vs $2^{nd}$ | 0.1942 | 0.3236667 | F |
| Control $1^{st}$ vs *Orai* RNAi $1^{st}$ | 0.8788 | 0.6660000 | F |
| Control $1^{st}$ vs *Stim* RNAi $1^{st}$ | 0.5328 | 0.8788000 | F |

**Figure 6E:** Chi-square comparing distribution of stretch sensitive cells

| | Chi-square value | DF | p-value |
|---|---|---|---|
| Control vs *Stim* RNAi | 33 | 4 | <0.0001 |
| Control vs *Orai* RNAi | 150.6 | 4 | <0.0001 |
| Stim RNAi vs *Orai* RNAi | 3.689 | 4 | 0.4497 |

**Figure 6G:** Chi-square comparing distribution of stretch sensitive cells

| | Chi-square value | DF | p-value |
|---|---|---|---|
| Control vs La3+ | 36.09 | 4 | <0.0001 |
| Control vs Store Depleted | 112.1 | 4 | <0.0001 |
| Store Depleted vs La3+ | 40.69 | 4 | <0.0001 |

**Figure 6J:** Fisher's exact test

| | p-value |
|---|---|
| control vs *GtACR* | 0.03216 |

**Figure 6K** Fisher's exact test with BH correction

| | p-value | q-value | Significance (q<0.1) |
|---|---|---|---|
| Control $1^{st}$ vs $2^{nd}$ | 0.00037 | 0.00148000 | T |
| *Stim OE* $1^{st}$ vs $2^{nd}$ | 0.02386 | 0.03181333 | T |
| Control $1^{st}$ vs *Stim OE* $1^{st}$ | 0.002115 | 0.00423000 | T |

**Figure 6L:** Fisher's exact test with BH correction

| | p-value | q-value | Significance |
|---|---|---|---|
| Control 25C 1st vs Control 25C 2nd | 5.425e-05 | 7.233333e-05 | T |
| Control 30C 1st vs Control 30C 2nd | 1.033e-06 | 2.066000e-06 | T |
| *Epi>shi* 25C 1st vs *Epi>shi* 25C 2nd | 4.912e-07 | 1.964800e-06 | T |
| *Epi>shi* 30C 1st vs *Epi>shi* 30C 2nd | 0.05076 | 5.076000e-02 | F |

**Figure 6S2A** Fisher's exact test with BH correction

| | p-value | q-value | significance |
|---|---|---|---|
| Control Trial 1 vs Trial 2 | 0.002438 | 0.004876 | T |
| Orai RNAi Trial 1 vs Trial 2 | 0.4076 | 0.4076 | F |

**Figure 6S2B** Fisher's exact test with BH correction

| | p-value | q-value | significance |
|---|---|---|---|
| Control Trial 1 vs Trial 2 | 0.00157 | 0.003140 | T |
| Task6 RNAi Trial 1 vs Trial 2 | 0.004795 | 0.004795 | T |

**Figure 6S2C** Wilcoxon Rank-Sum test

| | p-value | significance |
|---|---|---|
| Control vs $La^{3+}$ block | 0.0003377 | T |

**Figure 6S2E**: Kruskal-Wallis test followed by Wilcoxon rank sum test with BH correction

| | p-value | q-value | Significance (q<0.05) |
|---|---|---|---|
| Kruskal-Wallis test | 0.00001332 | | T |
| Control vs *Stim* RNAi | 0.0009161 | 0.001374150 | T |
| Control vs *Orai* RNAi | 0.000005094 | 0.000015282 | T |
| *Stim* RNAi vs *Orai* RNAi | 0.3759 | 0.375900000 | F |

**Figure 6S2F**: Kruskal-Wallis test followed by Wilcoxon rank sum test with BH correction

| | p-value | q-value | Significance (q<0.05) |
|---|---|---|---|
| Kruskal-Wallis test | 0.004105 | | T |
| Control vs *Stim* RNAi | 0.001296 | 0.002592 | T |
| Control vs *Orai* RNAi | 0.8294 | 0.829400 | F |

**Table S1. Larval behaviors scored in this study**

| Behavior | Description / Criteria |
|---|---|
| Locomotion | Continuous forward movement with smooth, regular peristalsis |
| Pausing | Break in locomotion, lasting less than 1 second |
| Freezing | Break in locomotion that persists for longer than 1 second |
| Hunching | Contraction along the anterior-posterior axis; no forward / backward motion or curling |
| Backward locomotion | Anterior-to-posterior peristalsis accompanied by movement in reverse |
| Writhing crawl | Forward movement with anterior and posterior sweeping of the body, tilting onto the side axis |
| C-bending | Bending into a c-shape, with both head and tail bending in the same direction; includes incomplete rolls |
| Rearing | Upwards bending or arching of anterior part of the body, with posterior part largely stationary |
| Rolling | Larva completes a full 360-degree rotation of its body |

**Table S2. Solutions used in calcium imaging studies.**

| **HL3.1, *Drosophila* saline** | | | |
|---|---|---|---|
| **Compound** | **M.W. (g/mol)** | **1X Concentration (mM)** | **For 1L (1X)** |
| NaCl | 58.44 | 120 | 7.013 g |
| KCl | 74.55 | 5 | 0.373 g |
| Proline | 115.13 | 5 | 0.576 g |
| HEPES | 238.30 | 10 | 2.383 g |
| Trehalose | 378.33 | 5 | 3.78 g |
| Sucrose | 342.30 | 32.5 | 11.125 g |
| CaCl2 | 1M stock | 1.5 | 1.5 ml from 1M stock |
| MgCl2 | 1M stock | 1 | 1 ml from 1M tock |
| | | pH (with NaOH) | 7.15 |
| | | Osmolarity (mOsm/L) | 310 |

| **Zero Ca2+ / EGTA HL3.1** | | | |
|---|---|---|---|
| **Compound** | **M.W. (g/mol)** | **1X Concentration (mM)** | **For 1L (1X)** |
| NaCl | 58.44 | 120 | 7.013 g |
| KCl | 74.55 | 5 | 0.373 g |
| Proline | 115.13 | 5 | 0.576 g |
| HEPES | 238.30 | 10 | 2.383 g |
| Trehalose | 378.33 | 5 | 3.78 g |
| Sucrose | 342.30 | 32.5 | 11.125 g |
| EGTA | 380.35 | 1.5 | 0.571 g |
| MgCl2 | 1M stock | 1 | 1 ml from 1M tock |
| | | pH (with NaOH) | 7.15 |
| | | Osmolarity (mOsm/L) | 310 |

| 20mM Ca2+ HL3.1 | | | |
|---|---|---|---|
| **Compound** | **M.W. (g/mol)** | **1X Concentration (mM)** | **For 1L (1X)** |
| NaCl | 58.44 | 101.5 | 5.932 g |
| KCl | 74.55 | 5 | 0.373 g |
| Proline | 115.13 | 5 | 0.576 g |
| HEPES | 238.30 | 10 | 2.383 g |
| Trehalose | 378.33 | 5 | 3.78 g |
| Sucrose | 342.30 | 32.5 | 11.125 g |
| CaCl2 | 1M stock | 20 | 20 ml from 1M stock |
| MgCl2 | 1M stock | 1 | 1 ml from 1M tock |
| | | pH (with NaOH) | 7.15 |
| | | Osmolarity (mOsm/L) | 310 |

| **Isotonic** (modified from *Drosophila* saline to maintain ionic balance across osmolarities) | | | |
|---|---|---|---|
| **Compound** | **M.W. (g/mol)** | **1X Concentration (mM)** | **For 1L (1X)** |
| NaCl | 58.44 | 90 | 5.260 g |
| KCl | 74.55 | 5 | 0.373 g |
| Proline | 115.13 | 5 | 0.576 g |
| HEPES | 238.30 | 10 | 2.383 g |
| Trehalose | 378.33 | 5 | 3.78 g |
| Sucrose | 342.30 | 92.5 | 31.663 g |
| CaCl2 | 1M stock | 1.5 | 1.5 ml from 1M stock |
| MgCl2 | 1M stock | 1 | 1 ml from 1M tock |
| | | pH (with NaOH) | 7.15 |
| | | Osmolarity (mOsm/L) | 310 |

| 85% Hypoosmotic | | | |
|---|---|---|---|
| **Compound** | **M.W. (g/mol)** | **1X Concentration (mM)** | **For 1L (1X)** |
| NaCl | 58.44 | 90 | 5.260 g |
| KCl | 74.55 | 5 | 0.373 g |
| Proline | 115.13 | 5 | 0.576 g |
| HEPES | 238.30 | 10 | 2.383 g |
| Trehalose | 378.33 | 5 | 3.78 g |
| Sucrose | 342.30 | 46.5 | 15.917 g |
| CaCl2 | 1M stock | 1.5 | 1.5 ml from 1M stock |
| MgCl2 | 1M stock | 1 | 1 ml from 1M tock |
| | | pH (with NaOH) | 7.15 |
| | | Osmolarity (mOsm/L) | |

| 70% Hypoosmotic | | | |
|---|---|---|---|
| **Compound** | **M.W. (g/mol)** | **1X Concentration (mM)** | **For 1L (1X)** |
| NaCl | 58.44 | 90 | 5.260 g |
| KCl | 74.55 | 5 | 0.373 g |
| Proline | 115.13 | 5 | 0.576 g |
| HEPES | 238.30 | 10 | 2.383 g |
| Trehalose | 378.33 | 5 | 3.78 g |
| Sucrose | 342.30 | 0 | 15.917 g |
| CaCl2 | 1M stock | 1.5 | 1.5 ml from 1M stock |
| MgCl2 | 1M stock | 1 | 1 ml from 1M tock |
| | | pH (with NaOH) | 7.15 |
| | | Osmolarity (mOsm/L) | |
